# Differences in sperm swimming speed and morphology between the three genetic morphs in the ruff sandpiper

**DOI:** 10.1101/2023.07.27.550846

**Authors:** Martin Bulla, Clemens Küpper, David B Lank, Jana Albrechtová, Jasmine L Loveland, Katrin Martin, Kim Teltscher, Margherita Cragnolini, Michael Lierz, Tomáš Albrecht, Wolfgang Forstmeier, Bart Kempenaers

## Abstract

**Background:** The ruff sandpiper *Calidris pugnax* is a Palearctic lekking shorebird with three genetic morphs determined by an autosomal inversion. Male morphs differ strikingly in body size, ornaments, endocrinology and mating behavior. Aggressive Independents represent the ancestral haplotype, while female-mimicking Faeders and semi-cooperative Satellites are the inverted haplotypes. Because one inversion breakpoint is homozygous lethal, the inverted haplotypes cannot recombine and are expected to accumulate deleterious mutations. The inversion regions also harbor genes involved in spermatogenesis. However, it remains unknown whether the genetic differences between the morphs also translate into differences in sperm traits. Here, we use a captive-bred population of ruffs to compare sperm swimming speed and morphology among the morphs.

**Results:** Ruff sperm morphologically resembled those of passerines, but moved differently, vibrating from side to side while slowly moving forward, rather than rotating while moving forward. Faeder sperm moved the slowest, which is consistent with the prediction of genetic deterioration over time. However, against expectation, sperm of Independents did not seem to be of the highest quality, i.e., their sperm were not the fastest nor the least variable, and they had sperm with the shortest tail and midpiece. Although the midpiece contains the energy-producing mitochondria, sperm midpiece length was not associated with sperm swimming speed. Instead, two of three velocity metrics weakly positively correlated with head length (absolute and relative).

**Conclusions:** The three genetically determined ruff morphs showed subtle differences in swimming speed and in the length of some sperm components. However, the between-morph differences in sperm swimming speed were not linked to the differences in morphology. We conclude that there is at best limited evidence for lower-quality sperm in the morphs that carry the inversion, and suggest that the potential for the evolution of morph-specific sperm adaptations may be limited in this system.

**Lay Summary:** The ruff sandpiper is a shorebird that exhibits three genetically distinct types of males, which differ markedly in body size, ornaments, hormones, and mating behavior. Aggressive Independents represent the type that evolved first. Semi-cooperative Satellites and female-mimicking Faeders evolved later through a specific type of genetic rearrangement called an ‘inversion’, in which a segment of a chromosome reversed in orientation. Due to the nature of this inversion, Satellite and Faeder chromosomes are expected to deteriorate over time. However, it remains unclear whether the genetic differences between these morphs, which affect physiological and behavioral traits, also translate into differences in sperm traits. We used a captive population of ruffs to compare sperm swimming speed and length measurements between the three types of males. Faeder sperm was the slowest, which is consistent with expectations based on genetic deterioration over time. However, against our expectations, the sperm of Independents does not appear to have better performance characteristics. Although the midpiece of a sperm is responsible for energy production, the length of the midpiece did not relate to sperm swimming speed.

## Background

In many animal species, males use different behavioral strategies to obtain fertilizations (Dougherty et al., 2022; Kustra & Alonzo, 2020; Mank, 2023). For example, dominant males may display, while other males “steal” copulations (Gross, 1996) by pretending to be a female or by positioning themselves in between the dominant male and the female that is about to copulate, a tactic referred to as “sneaking” (Jukema & Piersma, 2006; Shuster & Wade, 1991). Consequently, males that use alternative mating tactics often experience different levels of sperm competition, i.e., competition between the sperm from different males to fertilize one or more eggs (Parker, 1970). For example, if females prefer dominant males, it is likely that sneaker males more often experience sperm competition than dominant males. Comparative studies have shown that species experiencing higher levels of sperm competition typically produce sperm that swim faster, are longer (including longer sperm components), and less variable in length as a result of intense postcopulatory selection on optimal sperm length (Laskemoen et al., 2012; Lifjeld et al., 2010; Lipshutz et al., 2022; Lüpold et al., 2020; Simmons & Fitzpatrick, 2012). Within species, sperm fertilization ability has been linked to sperm viability (Gage et al., 2004; Garcia-Gonzalez & Simmons, 2005), swimming speed (Birkhead et al., 1999; Gage et al., 2004), and length (Bennison et al., 2014; Garcia-Gonzalez & Simmons, 2007; Lüpold et al., 2012). Thus, selection may have favored different sperm traits in males that consistently differ in mating behavior (Dougherty et al., 2022; Kustra & Alonzo, 2020). Despite the general expectation that sneaker males may have to produce more or more competitive sperm (higher quality or performance) to sire offspring, existing theory and data do not make clear predictions about how sperm traits of sneaker males should differ from those of other males (Dougherty et al., 2022; Kustra & Alonzo, 2020). Moreover, the evolutionary potential for sneakers to evolve superior sperm to compensate for their lower chances to copulate may be limited, because if sneaker males are rare in the population, the probability that a sneaker-specific beneficial mutation arises is low.

In the ruff *Calidris pugnax*, a lekking shorebird, males occur in three distinct morphs with striking differences in body size, ornaments, endocrinology and reproductive behavior (Hogan-Warburg, 1966; Höglund & Lundberg, 1989; Jukema & Piersma, 2006; Küpper et al., 2016; Loveland et al., 2021a; van Rhijn, 1991; Widemo, 1998). (1) Aggressive Independents make up 80-90% of all males and show a spectacular diversity of predominantly dark ornamental plumages. The Independents hold small display courts on a lek. Among these, the dominant male obtains the majority of matings (Vervoort & Kempenaers, 2019; Widemo, 1997). (2) Submissive and slightly smaller Satellites make up 10-20% of males, show predominantly white ornamental plumage, and co-display with court-holders on leks (Hogan-Warburg, 1966; Höglund & Lundberg, 1989; van Rhijn, 1991; Widemo, 1998). Although Satellites may “steal” some copulations, Independents may benefit from having Satellites on the lek because their presence helps attract females to the display court of that male (Hugie & Lank, 1997; Tolliver et al., 2023). (3) Faeders are rare (∼1%), mimic females in appearance (lack of ornamental plumage and smaller body size), and attempt to sneak copulations when a female solicits an ornamented displaying male (Jukema & Piersma, 2006). Despite striking differences in body size, the three morphs have similar testes size (Figure 1; Jukema & Piersma, 2006; Küpper et al., 2016), and hence presumably can produce similar numbers of sperm. However, the morphs differ in their reproductive endocrinology. During the lekking season, Independent males have higher levels of circulating testosterone, whereas Satellites and Faeders have higher levels of androstenedione, a testosterone precursor (Küpper et al., 2016; Loveland et al., 2021a).

**Figure 1:**
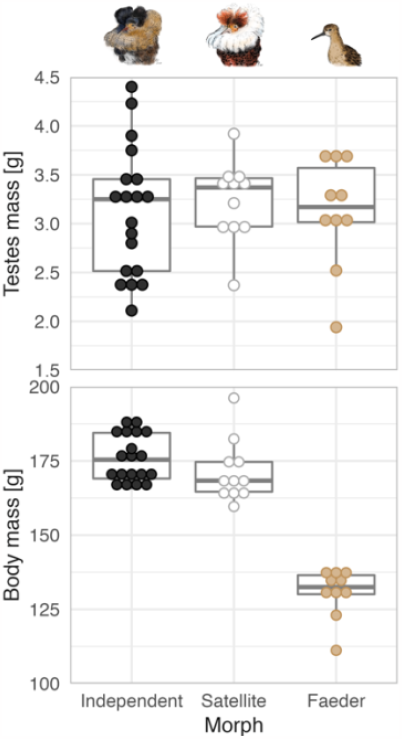
Testes mass and body mass in relation to morph in the ruff. Each dot represents an individual, dot color highlights the morph (black: Independent, white: Satellite, beige: Faeder). Boxplots depict the median (horizontal line inside the box), the 25^th^ and 75^th^ percentiles (box) and the 25^th^ and 75^th^ percentiles ±1.5 times the interquartile range or the minimum/maximum value, whichever is smaller (bars). Data from (Loveland et al., 2021b) and from an additional 5 males collected under the same protocol in 2022 (3 Independents, 1 Satellite and 1 Faeder; our unpublished data). Dots stacked using the ‘geom_dotplot’ ggplot2 R-function (Wickham, 2016). Ruff morph illustrations by Yifan Pei under [Creative Commons Attribution (CC BY 4.0)](https://creativecommons.org/licenses/by/4.0/).

Two observations suggest that sperm competition in ruff is intense. First, more than half of all clutches are sired by multiple males (Lank et al., 2002; Thuman & Griffith, 2005). Second, ruffs have the longest sperm measured among shorebirds (Johnson & Briskie, 1999). Female ruff visit one or multiple leks, and copulate during 3-5 days starting about 5 days before laying their first egg, and they produce a clutch of 3 or 4 eggs (Lank et al., 2002). As in other birds, females store sperm in sperm storage tubules prior to fertilization (Briskie & Montgomerie, 1993). Females of a given morph are approximately 40% smaller than males of the same morph (Giraldo-Deck et al., 2020; Lank et al., 2013). Presumably because of their smaller size, Faeder females have markedly lower reproductive success compared to Independent and Satellite females (Giraldo-Deck et al., 2022).

The phenotypic differences between the morphs are entirely genetically determined by a 4.5 Mb autosomal inversion region (Küpper et al., 2016; Lamichhaney et al., 2016). Faeders and Satellites carry unique inversion haplotypes. The Faeder haplotype arose first (about 4 million years ago), whereas the Satellite haplotype originated from a rare recombination event between the Independent and Faeder genotypes several hundred thousand generations later, about 70,000 years ago (Hill et al., 2023; Lamichhaney et al., 2016). Because one inversion breakpoint is homozygous lethal, the inversion haplotypes cannot recombine and thus only occur in hemizygous state (i.e. always combined with one Independent haplotype). As a result, the inversion haplotypes are expected to accumulate deleterious mutations. Although the genetic differences between the morphs affect numerous phenotypic traits, it is unknown whether they also affect sperm traits.

Intuitively, one might predict that the three morphs should differ in ejaculate size and in sperm traits due to differences in copulation opportunity and frequency. When a female visits a lek and solicits a copulation, the dominant Independent male is more likely to copulate once or multiple times with that female compared to a Satellite male (Lank et al., 2002). Although behavioral evidence is lacking, it is likely that copulation opportunities for Faeders are even more limited. If true, and given that the male morphs do not differ in testes size (Figure 1), sperm allocation theory predicts that Satellites and Faeders will transfer larger ejaculates per copulation than the dominant Independents (Pizzari et al., 2003). Differences in ejaculate size and in the presence of multiple ejaculates will produce different cloacal and oviductal environmental conditions in which sperm competition occurs. Such differences may have selected for different optimal sperm traits in the three morphs. Specifically, selection pressure on Faeders may be more intense as their ejaculate is more likely to compete with ejaculates of other males. Although an evolutionary scenario that leads to more competitive sperm in Faeders and Satellites may seem appealing, we argue that it is unlikely given the genetic make-up of the ruff morphs.

First, sperm traits and testicular gene expression may already have differed between morphs from the moment the morphs originated. (i) Two genes regulating spermatogenesis reside within the autosomal inversion that drives the Satellite and Faeder phenotypes (Küpper et al., 2016). The inversion contains the gene *GAS8*, which has been linked to sperm motility in mice, and the gene *SPATA2L*, a paralog of *SPATA2*, which is involved in spermatogenesis (Graziotto et al., 1999; Küpper et al., 2016; Onisto et al., 2000; Yeh et al., 2002). Moreover, expression of *SPATA2L* in testes of both Satellites and Faeders is biased toward the inversion allele (Loveland et al., 2021b). (ii) The inversion also influences expression of genes located outside of the inversion (Maney & Küpper, 2022). For example, *STAR*, a gene responsible for providing the cholesterol substrate required for sex hormone synthesis, is located outside of the inversion, but is overexpressed in the testes of Satellites and Faeders (Loveland et al., 2021a). (iii) Some genes or regulatory elements within the inversion region that have strong pleiotropic effects may also affect spermatogenesis.

Second, sperm trait differences between the morphs could also have arisen by *de novo* mutation after the origin of the three morphs, but through a different evolutionary scenario than the one described above. The genetic background outside of the inversion has been selected to function well in the context of the most abundant morph, i.e., the Independents (Giraldo-Deck et al., 2022). In contrast, even under strong selection due to intense sperm competition, Faeders, and to a lesser extent Satellites, have limited evolutionary potential due to (i) small population size, (ii) a small part (0.3%) of the genome within which a beneficial mutation would have to arise and (iii) a small number of genes within the inversion (only two of the ∼125 genes identified) have known effects on sperm performance or morphology (Küpper et al., 2016). Under these conditions, it seems less likely that evolution will have led to more competitive sperm in Faeders and Satellites. In fact, if anything, it seems more likely that the two inversion morphs will have sperm of inferior quality, because the inversions do not recombine and deleterious mutations can thus accumulate inside the inversion haplotypes (Berdan et al., 2021). Under this scenario, we predict that Faeders have the lowest quality sperm because, (i) a larger number of genes may have accumulated potentially harmful mutations in the Faeder haplotype, given that it arose several million years earlier than the rather recent Satellite haplotype (Hill et al., 2023; Küpper et al., 2016; Lamichhaney et al., 2016) and (ii) the recombination through which Satellites arose may have re-instated “good” copies of a gene that (directly or indirectly) affects sperm quality.

Here, we use sperm sampled from a captive breeding population of ruffs to address three issues. First, we assess differences in sperm swimming speed and sperm (component) length between the three morphs. If longer sperm is better at fertilization (see references above) and the sperm of inversion morphs has been disadvantaged due to mutation accumulation, we expect the sperm of Independents to be (1) faster swimming, (2) longer, and (3) less variable in length. We would also expect the sperm of Independents to have (4) a longer midpiece (energetic component; Cardullo & Baltz, 1991; Knief et al., 2017; Míčková et al., 2023), (5) a longer tail and flagellum (midpiece + tail, kinetic component; Cardullo & Baltz, 1991), and possibly (6) a longer midpiece and flagellum length relative to total sperm length (Humphries et al., 2008).

Second, we test whether variation in ruff sperm morphology predicts variation in sperm swimming speed. Although a positive relationship between overall sperm length, midpiece or flagellum length and swimming speed can be predicted, previous studies, including those in birds, have shown inconsistent results (reviewed in Bennison et al., 2016; Cramer et al., 2021; Fitzpatrick & Lüpold, 2014; Kim et al., 2017; Knief et al., 2017; Míčková et al., 2023; Støstad et al., 2018).

Third, we describe the motility and shape of ruff sperm in comparison with that of other bird species. Sperm motility has not yet been described in scolopacids and only rarely in other species of shorebirds, and sperm shape has only been sketchily described in a few species in this group (Johnson & Briskie, 1999; Retzius, 1909). Furthermore, scolopacid sperm superficially resemble the sperm of passerines, but differ from those of all other (so far evaluated) avian lineages (Retzius, 1909).

## Methods

We collected and analyzed the data based on [*a priori* designed protocols](https://raw.githack.com/MartinBulla/ruff_sperm_v2/main/Protocols/Preregistration_v3.html). For any deviations from the (see [Supplement](https://raw.githack.com/MartinBulla/ruff_sperm_v2/main/Supplement.pdf) Methods S0; Bulla et al., 2023).

### Captive population

A population of ∼300 ruffs is housed at the Max Planck Institute for Biological Intelligence, Seewiesen, Germany (for details see [Supplement](https://raw.githack.com/MartinBulla/ruff_sperm_v2/main/Supplement.pdf) Methods S1). The population was founded in 2018 with 23 ruffs obtained from Dutch breeders, 5 ruffs from German breeders and 194 ruffs from Simon Fraser University, Vancouver, Canada. The latter were individuals from a captive-bred population founded with 110 ruffs hatched from wild eggs collected in Finland in 1985, 1989 and 1990, plus two Faeder males brought from the Netherlands in 2006 (Lank et al., 2013; Lank et al., 1995). Each individual of the population has been genotyped for its morph using a set of six single-nucleotide polymorphism markers located in the inversion region (Giraldo-Deck et al., 2020).

### Sperm sampling

In May and June 2021, we collected sperm by abdominal massage (for a detailed protocol of sperm collection – including video - and sample preparation see https://s.gwdg.de/UlejLot) or by electro-stimulation (Lierz et al., 2013). The length (15 mm) and diameter (4 mm) of the electro-stimulation probe, as well as the electric current and the number of electric impulses was adapted to the sampled individuals. The probe was inserted into the urodeum of the cloaca to stimulate the ampullae ductus deferens using three 1 s electric current impulses. The voltage was increased gradually from 0.09 V to a maximum of 1.0 V until contractions of the cloaca and the muscles of the tail were observed; each impulse was followed by a 2-3 s break. In June, we only sampled sperm using abdominal massage. The morphs were sampled in a haphazard order to avoid a potential confounding effect of sampling order. In both months, we attempted to obtain sperm by abdominal massage from all males. We obtained at least one ejaculate sample from 92 males (59 Independents, 25 Satellites and 8 Faeders).

Ejaculates (∼0.5–3μl) were pipetted from the cloaca and immediately diluted and gently mixed in 50μl of preheated (40°C) Dulbecco’s Modified Eagle’s Medium (Advanced D-MEM, Invitrogen™). To record sperm swimming speed, we then pipetted an aliquot of 2.5μl onto a standard 20μm two-chamber count slide (Leja, The Netherlands) placed on a thermal plate (Tokai Hit, Tokai Hit Co., Ltd.) kept at 40 °C. When sperm densities on the slide were too high, we took a new aliquot and further diluted the sample. We then recorded sperm movements. By limiting the time between ejaculate collection and recording, we avoided that changes in sperm swimming speed over time (Cramer et al., 2016a) would influence the results. For morphology measurements, we pipetted an aliquot of 20μl into 50μl of a phosphate buffered saline solution (PBS; Sigma P-4417) containing 1% formalin.

### Measurement of sperm swimming speed

For each sperm sample we recorded sperm swimming speed at 25 frames per second for approximately 45s in eight different fields of the Leja slide under a 100x magnification, using phase contrast and a digital camera (UI-1540-C, Olympus) mounted on a microscope (CX41, Olympus) fitted with a thermal plate (Tokai Hit, Tokai Hit Co., Ltd.) kept at a constant temperature of 40°C. We confirmed that this temperature was appropriate based on cloacal temperature measurements of five female ruffs. For each female, we inserted a high-resolution temperature-probe connected to a MSR145 data logger (MSR Electronics GmbH, https://www.msr.ch/en/) into the cloaca, and logged the temperature every 5s for 3 min, i.e. until an asymptotic value was reached. Cloacal temperatures ranged between 40.6°C and 42.3°C, and are similar to the body temperatures measured in shorebirds (Charadriiformes; range: 40.4-41.8°C, N = 10 species) and in passerines (39.2-43.5°C, N = 16 species; (McNab, 1966).

A few weeks later, a single person (JA) analyzed each recorded field using the CEROS computer-assisted sperm analysis system (Hamilton Thorne Inc.). JA visually inspected the tracked objects, excluded non-sperm objects and static spermatozoa from the analysis (Cramer et al., 2016b; Laskemoen et al., 2010; Opatová et al., 2016), and noted the quality of the recording (e.g., presence of faeces). We recorded a median of 192 sperm cells per sample (mean = 206, range: 5 – 562; N = 134 recordings from a total of 92 males, 46 recorded in May and 88 in June) and for each sample the software estimated the mean curvilinear, straight-line, and average-path velocity (for details see [Supplement](https://raw.githack.com/MartinBulla/ruff_sperm_v2/main/Supplement.pdf) Methods S2 and Figure S1 and S2; for within-male seasonal repeatability see Figure S3 and S4, Table S1).

### Measurements of sperm morphology

Within 5 days of sample collection, we pipetted (i) 20μl of the PBS-1% formalin fixed sample into 50μl of 5% formalin for long-term storage and (ii) smeared 10μl onto a microscope slide, and let it dry at room temperature or on a heating block set at 37°C. The next day, each slide was rinsed gently with deionized water to remove salt crystals and dried at room temperature.

To each dried slide we added a drop of Bisbenzimide (Hoechst 33342; Molecular Probes) to stain the sperm nucleus, and a drop of MitoTracker™ Green FM Dye (Invitrogen™ M7514) to stain the sperm midpiece (mitochondria; Picture 1). Within 48h of staining, we inspected the microscope slides under 200x magnification with a Zeiss Axio Imager.M2 light microscope fitted with a Zeiss Axiocam 512 color camera (12 megapixel; 4250 × 2838, pixel size of 3.1μm × 3.1μm) and DAPI 465nm and green 519nm filters using Zeiss ZEN blue 3.1 imaging software. For each male we used a single sample (30 from May, 62 from June) and photographed at least 10 intact, normal-looking spermatozoa under 400x magnification (objective size 40, ocular size 10). We then selected the 10 best, single-sperm images per male for measurements. Previous studies also measured ten sperm per male to estimate the coefficient of variation in sperm traits (Kleven et al., 2008; Lifjeld et al., 2010). We then randomized and renamed all pictures, such that we measured sperm blind to the morph and the identity of the individuals.

**Picture 1:**
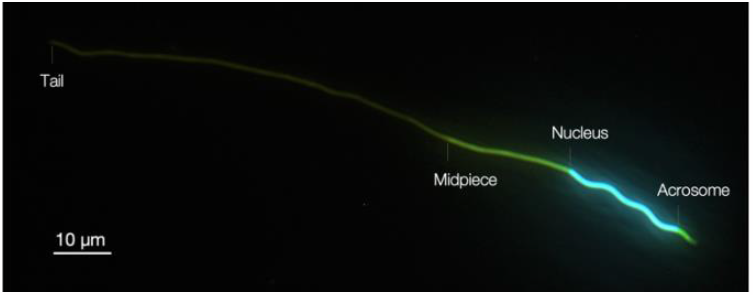
Stained ruff sperm. From left to right, faint greenish tail, green midpiece (mitochondria) stained by MitoTracker™ Green FM Dye, turquois nucleus stained by Hoechst 33342 and greenish acrosome.

For each sperm, we measured the length of the acrosome, the nucleus, the midpiece and the tail to the nearest 0.1μm using the open software Sperm Sizer 1.6.6 (McDiarmid et al., 2021; https://github.com/wyrli/sperm-sizer; for our Sperm Sizer protocol see https://s.gwdg.de/iGFz4F). To assess within- and between-observer repeatability of Sperm Sizer measurements, we selected 40 single-sperm pictures and remeasured them, once by the same, once by a different person. The measurements were highly repeatable, both within- and between persons (mean = 97%; Table S2). Nevertheless, all measurements used in the analyses were taken by the same person (KT). Each measured sperm and sperm part were numbered and referenced in the database and hence can be linked to the original picture as well as to the pictures of each part generated by Sperm Sizer, which contain the lines and measurement values (see Data in Bulla et al., 2023). This ensures transparency and allows re-measurement of the same sperm by the same or a different person.

We calculated (i) total sperm length as the sum of all parts, (ii) head length as the sum of acrosome and nucleus length, (iii) flagellum length as the sum of midpiece and tail length, (iv) relative midpiece length as midpiece length divided by total length and (v) relative flagellum length as flagellum length divided by total length. We then computed coefficients of variation within each male for each absolute trait as the standard deviation divided by the mean. In the [Supplement](https://raw.githack.com/MartinBulla/ruff_sperm_v2/main/Supplement.pdf), we report correlations between sperm components (Methods S3 and Figure S5), and estimates of within-male repeatability (Figure S3 and Table S1).

### Statistical analyses

All analyses were performed in R (R-Core-Team, 2022) using the ‘lm’ function to fit linear models and the ‘lmer’ function of the ‘lme4’ R-package to fit linear mixed-effects models (Gelman & Hill, 2007; Gelman & Su, 2021). We used the ‘sim’ function from the ‘arm’ R-package and a non-informative prior distribution (Gelman & Hill, 2007; Gelman & Su, 2021) to create a sample of 5,000 simulated values for each model parameter (i.e., posterior distribution). We report effect sizes and model predictions as medians, and the uncertainty of the estimates and predictions as Bayesian 95% credible intervals represented by the 2.5 and 97.5 percentiles (95% CI) from the posterior distribution of 5,000 simulated or predicted values. Unless stated differently, plots were created with ‘ggplot’ function from the ‘ggplo2’ R-package (Wickham, 2016).

To investigate whether sperm traits differ between the morphs we fitted linear models with each sperm trait as a response variable, and with male morph as a predictor (three-level factor). For analyses of velocity, we primarily used June values because the breeding season was at its peak in June and we obtained data from all but four males (for those four we used May values). We added the number of tracked sperm cells per sample (ln-transformed) as a covariate to the model to control for potential effects of sperm density or sample quality. For analyses of morphology, we used average male values. Alternative models using all velocity recordings, using individual sperm measurements, or controlling for sampling month or aviary gave similar results (Figure S6). Because correlations between sperm traits and inbreeding were weak (mean r = -0.09, range: -0.26 to 0.07; Figure S7, Methods S4), we did not include the inbreeding coefficient in the main models. Furthermore, the results were not confounded by relatedness (estimated based on microsatellite genotyping data) of the sampled individuals (see Methods S4, Table S3, Figure S8).

To investigate whether sperm morphology explained variation in sperm swimming speed, we fitted a set of linear models with each velocity measure as the dependent variable (using predominantly June values), and with the male average of each sperm morphology trait as a predictor. All models were controlled for the number of tracked sperm (ln-transformed) and morph (three-level factor). As the relationship between sperm swimming speed and morphology might be non-linear (Bennison et al., 2016), we also fitted a set of linear models that included the quadratic term of each morphology trait (2^nd^ order polynomial). We then tested whether the quadratic term improved the model fit, using Akaike’s Information Criterion corrected for sample size (Anderson, 2008), with the ‘AICc’ function from the ‘MuMIn’ R-package (Bartoń, 2022). Because this was not the case for any trait (Table S4), we report results from the simple model without the quadratic term. To investigate whether the effect of one sperm component (head, midpiece and tail length) was influenced by the other components, we also ran a multivariate model that contained the three sperm components as predictors. Correlations between the three predictors were low (Figure S5). The effect sizes and uncertainties for head, midpiece and tail length were similar in the multivariate and the univariate models (Figure S9) and we report the results from the univariate models in the main text.

## Results and Discussion

### Between-morph differences in sperm traits

Sperm swimming speed in vitro in a standard medium, measured as curvilinear velocity, did not differ between the three morphs (Figure 2 and [Supplement](https://raw.githack.com/MartinBulla/ruff_sperm_v2/main/Supplement.pdf) Figure S6 and Table S5; Bulla et al., 2023). However, when sperm swimming speed was measured as straight-line or average-path velocity, according to expectations based on mutation accumulation, Faeder sperm moved more slowly than the sperm of both Independents and Satellites, whereby the latter two morphs did not differ in sperm velocity (Figure 2 and S6, Table S5). If true, the slower sperm of Faeders appeared either with the origin of the Faeder genotype (i.e., the inversion) or later due to accumulation of deleterious mutations. In the latter case, recombination through which Satellites arose may have re-instated a “good” copy of a gene related to sperm velocity.

**Figure 2:**
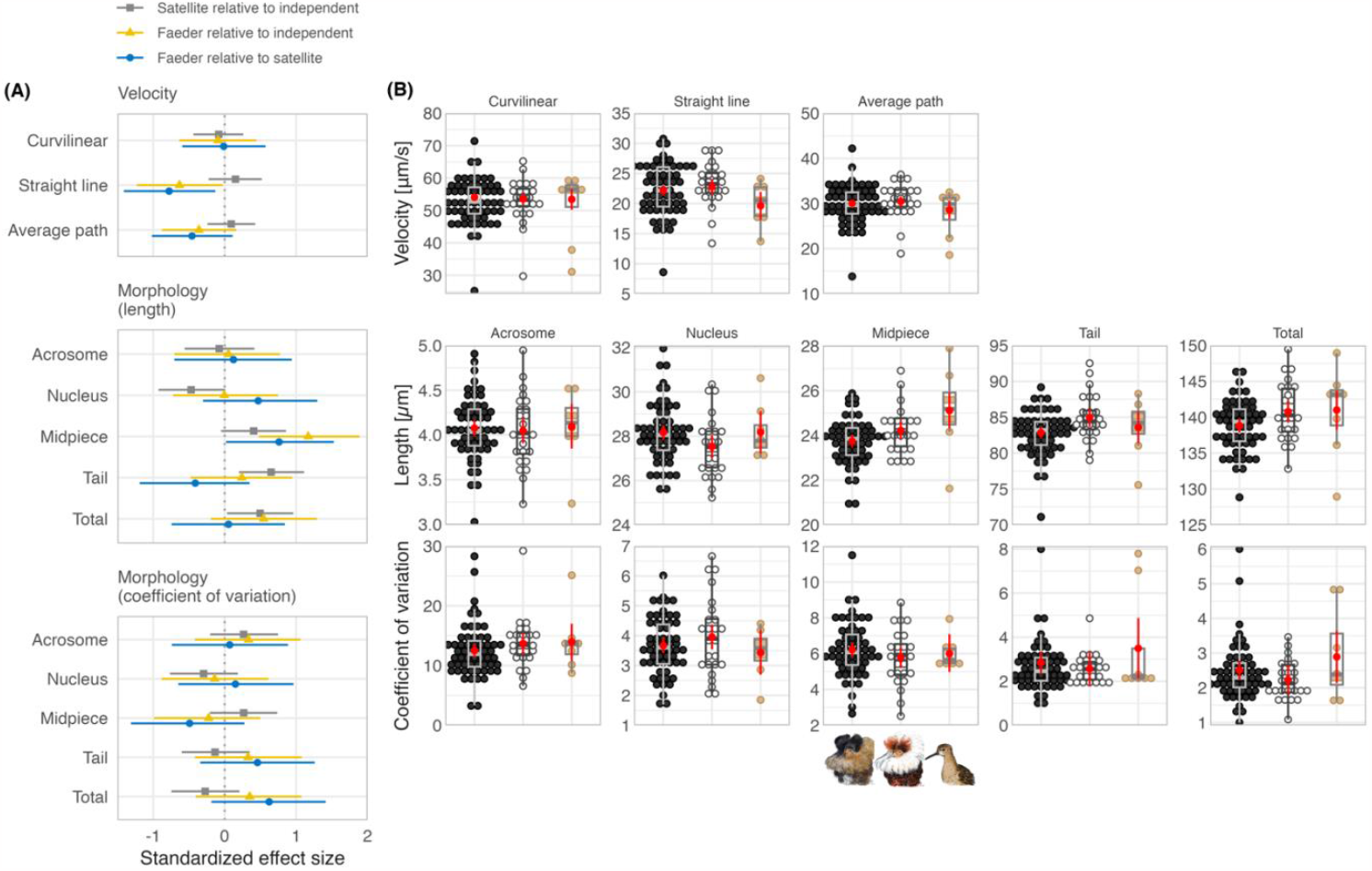
Differences in sperm traits between ruff morphs. (**a**) Model predictions of between-morph differences with their 95% CI (Table S5). (**b**) Morph-specific sperm traits, shown as individual data points with dot color highlighting morph (black: Independents, white: Satellites, beige: Faeders) and summarized as boxplots (see Fig. 1 for definition). To aid visualization, the outlier coefficients of variation for tail (18.7) and total length (9.8) in Independents are depicted as 8 and 6, respectivel y. The red dots with bars show model predictions and their 95% CI (Table S5). Dots were stacked using the ‘geom_dotplot’ ggplot2 R-function (Wickham, 2016). (**a, b**) Velocity represents June values (with the exception of four males with May values only). Morphology represents average trait length and coefficients of variation are based on measurements of 10 sperm cells per male. For estimates of head, flagellum (absolute and relative) and relative midpiece length, see Fig. S6 and S8. Ruff morph illustrations by Yifan Pei under [Creative Commons Attribution (CC BY 4.0)](https://creativecommons.org/licenses/by/4.0/).

Sperm morphology differed weakly between the three morphs, mostly by less than one standard deviation (Figure 2 and S6, Table S5). Sperm of Independents were the shortest with the shortest midpiece and tail, and hence flagellum (midpiece and tail length combined). Acrosome and nucleus length, and hence head length, did not differ between Faeders and Independents. Satellite sperm had the shortest nucleus and head, but the longest tail. The midpiece was longest in Faeders and 1.19 standard deviations longer than the midpiece from Independents (95%CI: 0.46 - 1.88; Figure 2, Table S5). Total sperm length and flagellum length were similar for Faeders and Satellites (Figure 2 and S6). Consequently, midpiece length relative to total sperm length was largest for Faeders, but similar for Satellites and Independents; flagellum length relative to total sperm length was largest in Satellites and similar in the other two morphs (Figure S6). The coefficients of variation in sperm traits were noisy with unclear trends and sperm of Independents was not less variable than that of the Faeders and Satellites (Figure 2 and S8). These findings do not support the prediction of mutation accumulation that sperm in Satellites and particularly in Faeders should be of lower quality. Specifically, when compared to Independents, the sperm of the two derived morphs were not shorter or more variable in terms of midpiece, tail, and flagellum. Thus, sperm of Independents seemed of similar or lower (not better) quality than sperm of the derived morphs (Figure 2), although what constitutes a high quality sperm, especially within species, is still debated (Dougherty et al., 2022).

The lack of major differences in sperm morphology among the three male morphs suggests that the two genes related to spermatogenesis that reside within the inversion have little impact on the measured sperm traits. Such finding might be unsurprising given that tens to hundreds of genes influence spermatogenesis (Linn et al., 2021). Furthermore, the lack of major differences in sperm traits accords with previous studies showing no clear and consistent differences in sperm traits between males that use alternative mating tactics (Dougherty et al., 2022; Kustra & Alonzo, 2020). Although one expects a steeper selection gradient for Faeder and Satellite males that have a lower probability to copulate (reviewed in Dougherty et al., 2022; Kustra & Alonzo, 2020), selection will act on all male morphs to optimize sperm traits. Moreover, independent of the strength of sexual selection, Faeder-beneficial mutations are about 30,000 times less likely to arise than Independent-beneficial mutations, given a Faeder population size of 1% and the inversion being 0.3% of the diploid genome (1/(0.01× 0.003)).

It remains unclear whether and how the success of Faeder males would compensate for the lower reproductive success of Faeder females (Giraldo-Deck et al., 2022). The measured sperm traits seem to give the Faeder males little advantage over Independent males. However, there is a strong mating skew with only few Independents obtaining most copulations on a given lek (Tolliver et al., 2023; Vervoort & Kempenaers, 2019; Widemo & Owens, 1995), such that the average reproductive success of an Independent will be low. Thus, the rare Faeder males might only need moderate mating success through sneaking, facilitated by their female mimicking appearance and behavior (Jukema & Piersma, 2006) to compensate. Possibly, Faeders do better, because morphs differ in other, non-measured, but fitness-relevant sperm traits. For example, the longer midpiece of Faeder sperm might correlate with longer sperm survival inside the female reproductive tract. However, this would need further study.

### Relationship between sperm swimming speed and morphology

Sperm morphology had only weak and unclear effects on variation in sperm swimming speed (Figure 3, S9 and S10, Table S6). Longer sperm, sperm with a longer midpiece (absolute and relative) or sperm with a longer flagellum did not swim faster (r varied between 0.05 and 0.18; Figure S10). The lack of an association between sperm midpiece length and swimming speed may have several explanations. First, our method to measure sperm velocity might not reflect the swimming speed of the sperm within the seminal fluid or within the female reproductive tract. Note, however, that the way the sperm moved did not change after we placed the sperm into a solution that contained cloacal fluid of a ruff female (details not shown). Also, our method is the same as the one used to measure swimming speed in passerine sperm, and we used the exact same equipment and observers as in some of the studies on passerines (e.g., Knief et al., 2017; Míčková et al., 2023; Opatová et al., 2016). Second, although the midpiece contains the mitochondria, midpiece volume rather than length might be relevant for sperm swimming speed (Cardullo & Baltz, 1991; Mendonca et al., 2018), but see Cramer et al. (2022), or midpiece length might be associated with other sperm performance parameters such as longevity (which we did not measure). Third, it remains unclear whether oxidative phosphorylation in the midpiece is the primary source of energy for moving sperm (Turner, 2003).

**Figure 3:**
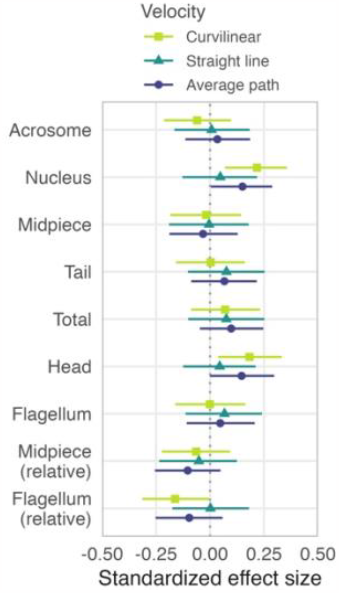
Effects of sperm morphology on sperm swimming speed. Shapes with horizontal bars represent estimated standardized effect sizes with their 95%CIs based on the joint posterior distribution of 5,000 simulated values generated from linear models, controlled for the number of tracked sperm (ln-transformed) and morph (Table S6). The results are based on single June-values for velocity (with the exception of four males with May-values only) and average trait length of 10 sperm cells per male. For raw data and predicted relationships see Figure S10.

Against the predictions (Humphries et al., 2008), sperm with longer heads (nucleus) or smaller relative flagellum length swam faster, when measured as curvilinear and average-path velocity (but not straight-line velocity), although these effects were small (< 0.25 of standard deviation, Figure 3; maximum r = 0.22, Figure S10). Theory predicts that sperm with a shorter head relative to flagellum should swim faster because a relatively shorter head reduces drag (Humphries et al., 2008). The few existing within-species studies provide equivocal evidence for this prediction (Cramer et al., 2015; Cramer et al., 2021; Helfenstein et al., 2009; Mossman et al., 2009), but the prediction has been supported in a between-species study in mammals (Tourmente et al., 2011) and in passerine birds (Lüpold et al., 2009). A recent study on passerines showed that sperm with a long acrosome, a short nucleus, a wide helical membrane, and a more pronounced waveform along the sperm head “core” swam faster (Støstad et al., 2018).

### Sperm motility and sperm shape in ruffs compared to other taxa

We observed that ruff sperm seemed to move in a different manner to that previously described for passerine and likely also for non-passerine sperm (video examples). Specifically, ruff sperm seem to ‘vibrate’ from side to side while slowly moving forward, whereas passerine sperm typically swim straight by rapidly rotating around their longitudinal axis (Ballowitz, 1888; Humphreys, 1972; Vernon & Woolley, 1999). We can exclude that the unusual head movements of ruff sperm were caused by a lack of depth in the standard 20μm two-chamber count Leja-slide, because using a deeper Leja-slide (100μm) resulted in similar movements (not recorded). However, our observations need further verification using other methods (Vernon & Woolley, 1999). If confirmed, the difference in movement may not be due to differences in general morphology, because ruff sperm have a helical, screw-like head (Picture 1), similar to that observed in passerine sperm (Retzius, 1909). Previous studies described that non-passerine sperm moves with regular helical (dextral) waves (Ballowitz, 1888; Humphreys, 1972; Vernon & Woolley, 1999) or via counter-clockwise turning of the entire sperm body (Bird & Laguë, 1977). Given the few descriptions and the lack of video recordings of non-passerine sperm movements (e.g., Cheng et al., 2002; Denk et al., 2005; Dogliero et al., 2017; Fischer et al., 2014; Gloria et al., 2014), it remains to be seen whether the way ruff sperm propel is unique to this species, or typical for other Scolopacids, shorebirds or Charadriiformes in general. Clearly, the directional movement of ruff sperm is distinct from the smooth, snake-like, movements of sperm of non-avian taxa (e.g., human or sea-urchin *Psammechinus miliaris* (Cosson et al., 2015; Gillies et al., 2009; Gray, 1955; Saggiorato et al., 2017; Smith et al., 2009), but see squid sperm (Bishop, 1958)).

The screw-like head of ruff sperm is similar to the head shape reported for other Scolopaci (sandpipers), but differs from the straight heads observed in other Charadriiformes, e.g., genus *Larus* or *Vanellus* (Retzius, 1909). Our results confirm that ruff sperm is remarkably long (median and mean = 139.5 µm, range: 105.7 - 150 µm) compared to other shorebirds, with species means ranging between 57 and 100 µm (Johnson & Briskie, 1999).

## Conclusions

Using sperm collected from a population of captive ruffs, we found between-morph differences in sperm swimming speed and length measures. As expected from mutation accumulation in the inversion, sperm of Faeders were the slowest, but – against expectation – sperm of Independents were not the fastest, longest or least variable. Furthermore, we found that sperm of ruffs moved differently than sperm of passerines, despite having a similar head shape (Ballowitz, 1888; Humphreys, 1972; Retzius, 1909; Vernon & Woolley, 1999). Our study shows at best weak associations between sperm swimming speed and length measurements that are inconsistent with general expectations. Our results corroborate other comparative work that revealed a lack of clear and consistent differences in sperm traits of males using alternative mating tactics (Dougherty et al., 2022; Kustra & Alonzo, 2020). Our study leaves open the question about whether and how Faeder males can achieve higher reproductive success to compensate for the lower reproductive output of Faeder females (Giraldo-Deck et al., 2022), but emphasizes little potential for the evolution of strong morph-specific sperm adaptations in this system.

## Availability of data and materials

All data, including sperm recordings and pictures, and computer code used to generate the results and Supplement (figures, tables and documents) of this study are freely available at GitHub: https://github.com/MartinBulla/ruff_sperm_v2 (Bulla et al., 2023).

## Acknowledgements

We thank the many people who took care of the captive ruffs, Melanie Schneider for genotyping, Callum S. McDiarmid for help with his Sperm Sizer software, Esteban Botero-Delgadillo for the script on homozygosity by locus, Jarrod Hadfield for advice on a genomic relationship matrix, Mihai Valcu for advice on statistics, Yifan Pei for ruff illustrations, and two anonymous reviewers for the comments.

## Author contributions

MB initiated the project with support of BK and DL. BK secured funding. MB, BK, TA, DL, CK, JL conceptualized the project. JL collected and measured the testes. MB, KT, TA, ML collected the sperm samples. JA measured velocity. KT and MB with support of TA optimized the photographing of the sperm. MB, MC and KT developed and tested the sperm measuring protocol. KT and MC prepared sperm samples, took sperm pictures and KT measured sperm morphology. MB curated and analyzed the data with input from BK, WF and TA. MB wrote the first draft and revised it with the help of BK. All authors commented and MB with BK, WF, CK and DL finalized the paper.

## Ethics declarations

### Ethics approval and consent to participate

The data were collected under license (311.5-5682.1/1-2014-020) of the Landratsamt Starnberg and in accordance with [Directive 2010/63/EU](https://eur-lex.europa.eu/LexUriServ/LexUriServ.do?uri=OJ:L:2010:276:0033:0079:EN:PDF).

### Consent for publication

Not applicable.

